# The feature-weighted receptive field: an interpretable encoding model for complex feature spaces

**DOI:** 10.1101/126318

**Authors:** Ghislain St-Yves, Thomas Naselaris

**Affiliations:** Medical University of South Carolina, Charleston, SC, USA

**Keywords:** Feature-weighted receptive field, Voxel-wise encoding model, Deep neural network, Visual cortex, fMRI

## Abstract

We introduce the feature-weighted receptive field (fwRF), an encoding model designed to balance expressiveness, interpretability and scalability. The fwRF is organized around the notion of a feature map—a transformation of visual stimuli into visual features that preserves the topology of visual space (but not necessarily the native resolution of the stimulus). The key assumption of the fwRF model is that activity in each voxel encodes variation in a spatially localized region across multiple feature maps. This region is fixed for all feature maps; however, the contribution of each feature map to voxel activity is weighted. Thus, the model has two separable sets of parameters: “where” parameters that characterize the location and extent of pooling over visual features, and “what” parameters that characterize tuning to visual features. The “where” parameters are analogous to classical receptive fields, while “what” parameters are analogous to classical tuning functions. By treating these as separable parameters, the fwRF model complexity is independent of the resolution of the underlying feature maps. This makes it possible to estimate models with thousands of high-resolution feature maps from relatively small amounts of data. Once a fwRF model has been estimated from data, spatial pooling and feature tuning can be read-off directly with no (or very little) additional post-processing or in-silico experimentation.

We describe an optimization algorithm for estimating fwRF models from data acquired during standard visual neuroimaging experiments. We then demonstrate the model’s application to two distinct sets of features: Gabor wavelets and features supplied by a deep convolutional neural network. We show that when Gabor feature maps are used, the fwRF model recovers receptive fields and spatial frequency tuning functions consistent with known organizational principles of the visual cortex. We also show that a fwRF model can be used to regress entire deep convolutional networks against brain activity. The ability to use whole networks in a single encoding model yields state-of-the-art prediction accuracy. Our results suggest a wide variety of uses for the feature-weighted receptive field model, from retinotopic mapping with natural scenes, to regressing the activities of whole deep neural networks onto measured brain activity.

## 1. Introduction

This paper describes and demonstrates the feature-weighted receptive field (fwRF) model, a new approach to building encoding models for visual brain areas. We initially developed the fwRF as a method for linking the visual features learned by deep artificial neural networks (DNNs) to activity in the human brain (measured, in our case, with fMRI). In recent years DNNs have been trained to perform visual processing tasks (e.g., human-level object recognition, natural image captioning, etc.) that previously could only be performed by biological visual systems [1]. They have also been shown to provide excellent models of processing in visual cortex [2, 3, 4]. For this reason, the internal representations used by DNNs provide a natural and compelling set of hypotheses about the visual features encoded by activity in real brains.

In previous work [5], we have argued that an excellent method for testing if a set of visual features (such as those learned by DNNs) is encoded by activity in the brain is to embed those features in an *encoding model*. An encoding model specifies a mapping from a set of visual features to a prediction of brain activity. The visual features in the model can be regarded as hypotheses about the visual features that might be encoded in brain activity. In encoding models, distinct visual features are each assigned a weight that indicates the importance of the visual feature for explaining measured brain activity. Important features will typically have large weights while unimportant features will have small weights. The weights for visual features are learned from a set of training data using an appropriate optimization algorithm—typically some form of regularized regression. Once the model weights have been learned, the model can be validated and compared to other models by testing its ability to predict brain activity in response to stimuli or task conditions that were not part of the training set.

The typical DNN is a gargantuan construct consisting of hundreds of thousands of nodes. The scale of these networks makes it challenging to fit them into the encoding model framework. In developing the fwRF we considered four specific challenges that together make up the performance goals of the fwRF model:

- *Expressiveness*: One way to fit DNNs into the encoding model framework is to regress each layer of a DNN onto brain activity independently. Although this approach has proven highly effective [2, 4], this method for reducing model scale comes at a cost of model expressiveness. We expect the costs to be severe in densely connected networks that do not admit an obvious decomposition in relatively small stacks of feature maps. Thus, an explicit performance goal of the fwRF modeling approach is to be able to construct voxelwise encoding models that simultaneously regress all feature maps in a DNN onto brain activity.
- *Interpretability*: Models with many thousands (or millions) of feature weights pose an obvious interpretive challenge. One aspect of this challenge is to extract simple visual cortical functional descriptors such as receptive field sizes and centers and tuning functions for the visual features in the model. Thus we sought a modeling approach that would yield explicit receptive field-like locations and pooling size descriptors, as well as explicit feature tuning functions that could be easily read-off from model parameters.
- *Scalability*: The regularized regression techniques we have used in the past [6, 7, 8] are likely to incur prohibitive memory or compute-time costs when scaled up to encoding models that use entire DNNs. Thus we sought to impose a set of reasonable constraints on our encoding model that would limit the growth of model parameters with model expressiveness without compromising the accuracy of model predictions.
- *Compatibility*: Any approach to building encoding models for visual areas should be able to use visual features from any source including, but not limited to, DNNs. Thus, while achieving the three goals mentioned above, we wanted a modeling approach that could also be used to construct encoding models that have already been proven to be effective, such as the Gabor wavelet pyramid model [6], the population receptive field model [9], the motion energy model [8], and models that use abstract features derived from other machine learning approaches [10].

To meet these performance goals we based the fwRF modeling approach on three key design principles:

- *Visual features of the model organized into feature maps*: In a fwRF model, visual features must be configured as pixels in a stack of feature maps. A feature map is a transformation of visual stimuli into abstract visual features. Each visual feature is a pixel in an image that preserves the topology of visual space (but not necessarily the native resolution of the stimulus). Note that this is a very general requirement, since features that consist of a single value (e.g., the nodes in a full connected layer of a DNN) can be treated as feature maps with a single pixel.
- *Explicit receptive field-like model*: Within visual areas, population activity at each point in the cortical sheet encodes visual features within a limited and contiguous region of the visual field. For this reason, the fwRF model contains an explicit receptive field-like model which we call the *feature pooling field* because it pools over pixels in feature maps (as opposed to pixels in the stimulus). The feature pooling field has an explicit center that indicates the location of a feature map that makes the largest contribution to the activity measured in the voxel, and a feature pooling radius that measures how quickly the contribution decays with distance from the center.
- *Space-feature separability*: In a fwRF model the location and radius of the feature pooling field are independent of the content of feature maps. The model thus assumes that activity measured in a single voxel will not encode distinct features at distinct locations, but rather a weighted combination of features at a single location. This *space-feature separability* makes the fwRF model scalable by keeping the complexity of the model independent of the resolution of the feature maps.

Here we show that the fwRF model can recover the location and radius of feature pooling fields and feature tuning functions that are consistent with known principles of cortical organization with minimal post-training analysis (interpretability), using either a small set of Gabor maps (compatibility) or a massive set of feature maps from a deep neural network (expressiveness), while achieving state-of-the art prediction accuracy within a reasonable (i.e., a few hours) training time (scalability).

## 2. Methods and materials

### 2.1. Data

The data used in this study are described in detail in Ref. [11]. Briefly, functional BOLD activity was measured in the occipital lobe with 4T INOVA MR scanner (Varian, Inc.) at a spatial resolution of 2mm × 2mm × 2.5mm and a temporal resolution of 1 Hz. During the acquisition, subjects viewed sequences of 20^*o*^ × 20^*o*^ greyscale natural photographs while fixating on a central white square. Photographs were presented for 1s with a delay of 3s between successive photographs. The data, available online at https://crcns.org/data-sets/vc/vim-1, is partitioned into distinct training and validation sets. The training set consists of estimated voxel activation in response to 1,750 photographs while the validation set consists of estimated voxel activation in response to 120 photographs. Figures in this study refer to data from subject 1 of the vim-1 dataset (similar results were obtained for subject 2).

### 2.2. Feature-weighted receptive field model

In this section we describe the general form of the feature-weighted receptive field (fwRF) model, the algorithm for optimizing the model to predict activity in individual voxels, two specific variants of the fwRF model and an alternative to the fwRF model.

#### 2.2.1. General form and motivation of the model

The fwRF model for a single voxel has three main components: a stack of feature maps, a vector of feature weights, and a feature pooling field.

Feature maps are maps of visual features over visual space (see Fig. 1A and B for examples). A feature map can be thought of as an image in which the pixels do not necessarily indicate the amount of light or color at a particular location, but may instead indicate the degree to which a potentially abstract visual feature is present or absent. A visual feature can be any sort of visual description of an image. Gabor wavelets are examples of visual features. Convolving an image with a vertically oriented Gabor filter, for example, produces a map of vertically oriented edge features. Objects can also be considered visual features. For example, we could obtain a “car” feature map by simply setting to 1 the value of image pixels occupied by a car and setting to 0 the value of all other pixels. Features can be much simpler than edges and/or objects. In the case of the population receptive field (pRF)model [9], the feature map is simply a binary map of the pixels occupied by a high-contrast stimulus (e.g., a wedge, ring, or bar).

**Figure 1:**
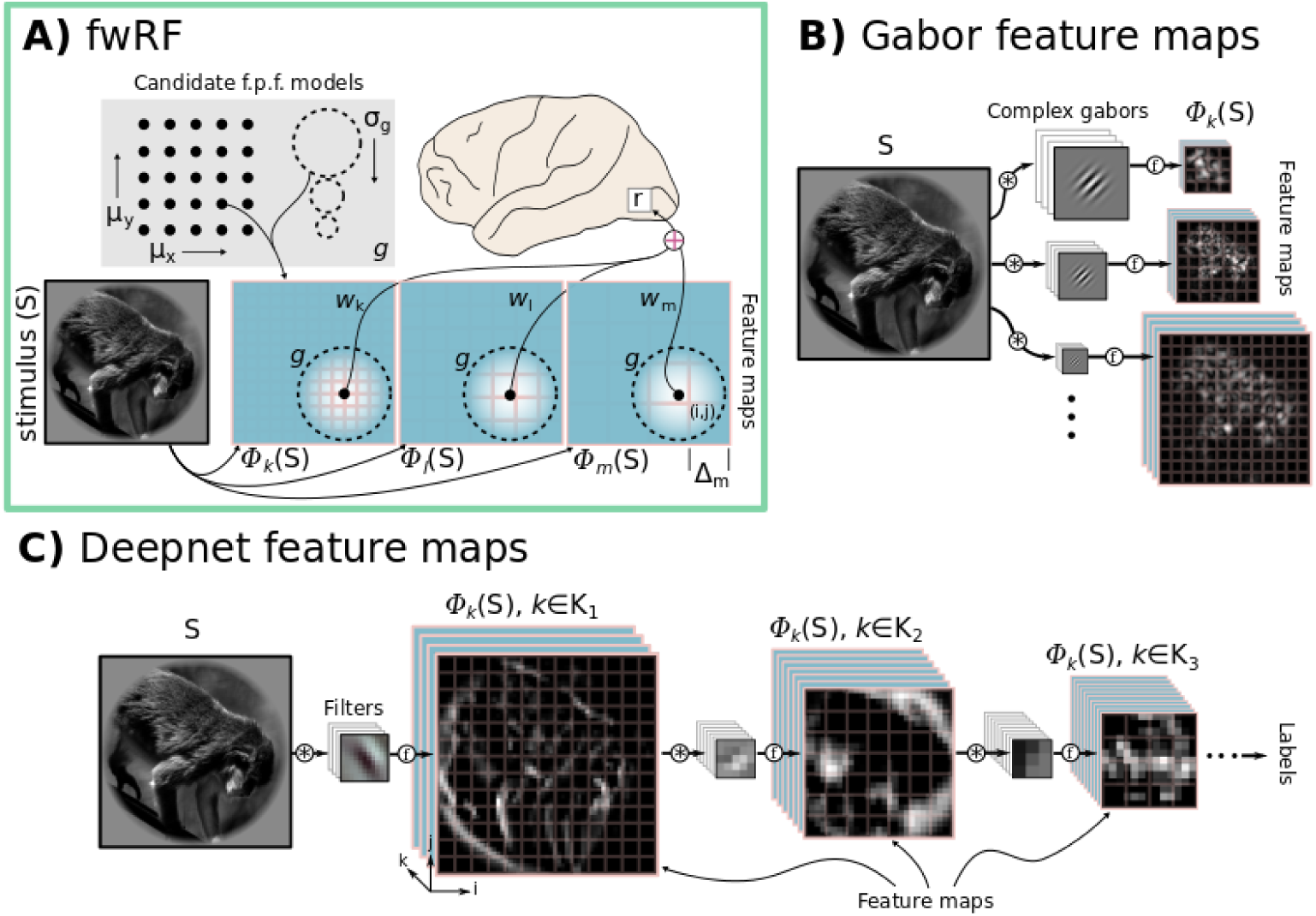
The fwRF model. (A) A schematic illustration of a fwRF for a single voxel (grey box on brain, top right). The fwRF predicts the brain activity measured in the voxel, *r*, in response to any visual stimulus, *S* (bottom left). The stimulus is transformed into one or more feature maps (three feature maps, Φ_k_, Φ_l_, and Φ_m_, are shown in blue with pink borders). The choice of feature maps is entirely up to the user, and reflects her hypotheses about the visual features that are relevant to brain regions of interest. The resolution of the feature maps (Δ, indicated by pink grids) can vary, although each feature map spans the same degree of visual angle as the stimulus *S*. Each feature map is filtered by a 2D Gaussian feature pooling field, *g*, that is sampled from a grid of candidate feature pooling fields (grey box at top left; candidate feature pooling field centers (*μ*_*x*_, *μ_y_*) are illustrated by the grid of black points, while candidate feature pooling field radii (*σ_g_*) are illustrated by dashed circles). The feature pooling field radius and location are the same for each feature map. The output of the feature pooling field filtering operation (illustrated as black dots in the center of the dashed feature pooling fields on each feature map) for each feature map is then weighted by a feature weight (black curves labeled *w_k_, w_l_, w_m_*). These weighted outputs are summed to produce a prediction of the activity *r*. In the text we describe an algorithm for selecting the optimal feature pooling field and feature weights for each voxel. (B) Gabor wavelet feature maps are constructed by convolving the input images with complex Gabor wavelets followed by a compressive nonlinearity (see text for details). (C) Deepnet feature maps were extracted the layers (labeled *K_i_*) of a deep convolutional network pre-trained to label images according to object category.

Formally, a feature map is a matrix function. Given an image *S_t_*, the feature map Φ_*k*_(*S_t_*) outputs a matrix where elements 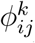 (*S_t_*) are feature map pixels. A fwRF model may include multiple (typically between 1 and 10^4^) feature maps, so we index each feature map by k, and let Φ = {Φ_*k*_(*S*)} denote the full set of *K* feature maps in a fwRF model.

Why should a fwRF model include more than one feature map? In many cases, we will not know what features will best explain activity in the brain regions of interest (ROIs). Thus, Φ will ideally include enough feature maps to capture the breadth of reasonable hypotheses about what is encoded in the activity of ROIs. We then use training samples (activity/image pairs) to infer which of the features are most important for explaining the activity of each voxel. We do this by assigning to each feature map in a fwRF model an associated *feature weight, w_k_*, that indicates how important the *k*^th^ feature map is for predicting how activity will vary across images. These feature weights are the “what” parameters of the model since they indicate the visual features that are important for explaining activity in the voxel. Note that when constructing multiple voxelwise encoding models, the feature maps Φ are the same for each voxel, but the *feature weights* vary across voxels.

A final and critical assumption of the fwRF model is that activity measured in individual voxels is driven most strongly by feature map pixels that are close to the center of the voxel’s feature pooling field. The farther the feature map pixels are from this center, the weaker the contribution they will make to measured brain activity. We assume that both the center and radius (in units of deg.) of the feature pooling field is the same across all feature maps. The feature pooling field can thus be thought of as a window on the feature maps that does not change from one map to the next. This is a reasonable assumption when the number of pixels in a feature map times the pooling size of the individual pixels is roughly constant across all feature maps. In such a case, the feature pooling field covers less and less pixels in a feature map as the resolution decreases, but each feature unit in turns inherently pools over a larger visual area.

In this treatment, we model the feature pooling field as an isotropic 2D Gaussian blob (although they could be more complicated functions)

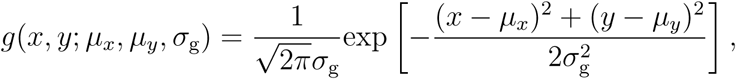
where the mean parameter ***μ*** = (*μ_x_, μ_y_*) is the feature pooling field center and the variance parameter *σ_g_* is the feature pooling field radius. To predict the response of the neuron or voxel to an image *S_t_*, the feature maps for that image are formed, the feature pooling field is applied to each feature map, and the feature weights are applied to these outputs. Formally:

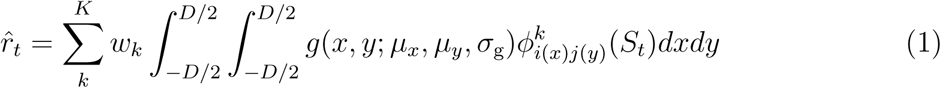

Where 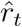 is the predicted activity in response to image *S_t_* and *D* is the total visual angle sustained by the image *S_t_*. The discretization depends on the resolution of the feature map under consideration such that 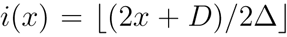 (likewise for j(y)) where Δ = *D/n_k_* is the visual angle sustained by one pixel of a feature map with resolution *n_k_* × *n_k_*. This definition reduces to a discrete weighted sum when the resolution of a feature map is very high relative to the size of the feature pooling field (i.e. when Δ ≪ *σ_g_*)and reduces to a single feature map spatial unit being exclusively selected when the size of the feature pooling field is smaller the resolution of the feature map. In practice, there is often an additional voxel-wise bias parameter *b*, which we omit for simplicity since it does not play a role in the validation accuracy.

When optimizing the model for a particular voxel, the goal is to infer values for the feature pooling field parameters *σ_g_* and ***μ***, and feature weights **w** = (*w_1_*, …, *w_K_*) that result in accurate predictions of the unit’s response to any image.

#### 2.2.2. Optimization algorithm for the fwRF model

The optimal fwRF model parameters were estimated by minimizing a least-squares cost for each voxel. Let us write the parameters concisely as Θ = (**w**, ***μ***, *σ*_g_), then the cost for a single voxel is

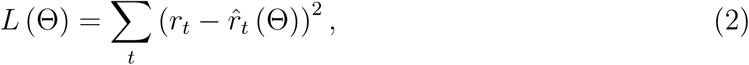
where *r_t_* is measured activity in response to image *S_t_*, and 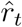 is the prediction of the fwRF model (as defined above) expressed as an explicit function of the model parameters Θ.

For fixed feature pooling field parameters, the fwRF model is linear in the feature weights w and one can optimize the feature weights via regularized regression. However, the model has a nonlinear dependence on the feature pooling field parameters ***μ*** and *σ_g_*. Therefore, we construct a grid of *candidate* feature pooling field locations and sizes. For each of these *G* candidate feature pooling fields, we perform stochastic gradient descent (with early stopping) on the feature weights. Under this procedure, for each candidate feature pooling field, the gradient of *L* (Θ) with respect to **w** is computed using 80% of the data samples in the training set. Gradient descent is performed for all candidate feature pooling fields for a fixed number of iterations, resulting in *G* candidate models for the voxel. The model that minimizes cost on the remaining 20% of the training data is considered the optimal model for the voxel and retained for further analysis. See the supplementary materials for a pseudocode detailing the optimization procedure.

#### 2.2.3. Software implementation

The fwRF models described here were implemented using Theano, a Python toolkit for machine learning [12]. A more detailed description of the main procedures is given in the supplementary materials (Algorithm A.1) as well as an estimate of the model scaling under various conditions (Figure A.1). Furthermore, executable IPython notebooks that illustrate the construction and training of the fwRF models described here are available online at https://github.com/styvesg/fwrf.

### 2.3. Details of the models

In the following, we will consider three different models: A fwRF model based on Gabor wavelets (Fig 1B), a fwRF model based on feature maps from a DNN (Fig 1C), and a deep network layerwise regression based on the same network (Fig 2).

**Figure 2:**
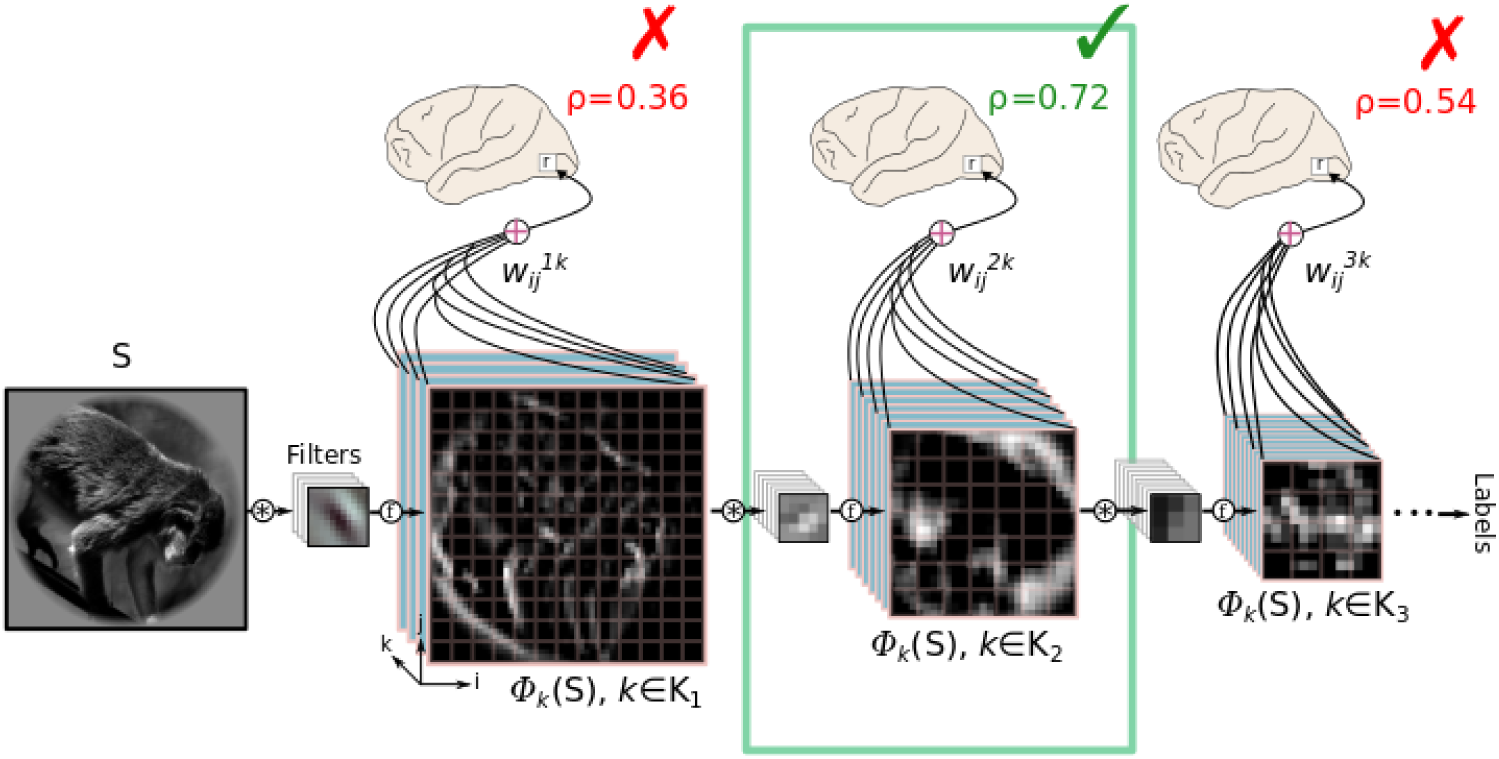
Deepnet layerwise ridge regression method. In the layerwise model feature maps (*bottom*) are supplied by the same DNN used in the Deepnet-fwRF model. In contrast to the fwRF method, the layerwise ridge regression has one feature weight (curved black lines labeled 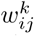) per feature map pixel (illustrated by pink grid overlaid onto feature maps). Feature weights are fit independently for each layer, resulting in *L* (where *L* is the number of layers in the DNN) distinct encoding models for each voxel. For each voxel (small box labeled *r* in the brain diagrams), the model with the best prediction accuracy (*ρ*) on a held-out model selection set is selected and retained for further analysis. In this illustration the model associated with the second layer of the DNN is selected (green box and checkmark) while the models for the first and third layers are discarded (red *X*’s).

#### 2.3.1. fwRF model with Gabor wavelet feature maps (Gabor-fwRF)

For the Gabor-fwRF model, each of the *K* feature maps is associated with a single 2*D* complex-valued Gabor wavelet. We’ll denote the gabor wavelet *h* (*ω*_k_, *Θ_k_*), where *ω_k_* and *Θ_k_* are the spatial frequency (cycles/degree) and orientation (radians) of the wavelet,respectively. Then for the Gabor-fwRF, 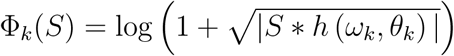 In this study,we used *K* = 96 gabors at 12 log-spaced spatial frequencies between 0.25 cyc./deg. and 6.0 cyc./deg. For each frequency we sampled 8 evenly-spaced orientations between 0 and 7π/8. The grid of candidate feature pooling fields included 16 radii between *σ_g_* = 0.25 and *σ_g_*= 8. Candidate feature pooling field centers were spaced 0.61 degrees apart (regardless of radius) for a total of *G* = 16, 384 candidate feature pooling fields. The model for each voxel was run for 20 epochs of stochastic gradient descent with batch size of 200 and step size of 10^−3^ starting from an initial state of **w** = **0**. Increasing the maximum number of epochs seemed to confer no further significant improvement.

#### 2.3.2. fwRF model with Deep Convolutional Neural Network feature maps (Deepnet-fwRF)

Each of the *K* feature maps in the Deepnet-fwRF model is associated with a feature map in one layer of a deep convolutional neural network. The network contains one input layer, 5 convolutional layers (interleaved with max-pool layers) and 3 fully-connected layers. The depth (number of feature maps) and resolution (square root of the number of pixels in each feature map) for each convolutional layer was (96, 55), (256, 27), (384, 13), (384, 13), (256, 13) respectively. The fully-connected layers contained 4, 096, 4, 096 and 1, 000 units respectively. The network was trained to classify images in the ImageNet database (http://image-net.org) according to 1, 000 distinct class labels. For a complete description of the network architecture and training, see [13]. The exact structure and trained network weights can be downloaded as part of the Caffe package [14] and is referred therein as the *bvlc_reference_caffenet* network (details can be found at http://caffe.berkeleyvision.org/model_zoo.html).

All feature maps from all convolutional layers, as well as up to 1024 units from the fully-connected layers (when the number of feature maps in a layer exceeded this amount, we used the first 1024 feature maps which exhibited the most variance to the training set), were used for the Deepnet-fwRF model resulting in a model with 4, 424 feature weights. The grid of candidate receptive fields included 10 receptive field sizes between *σ_g_* = 0.7 and *σ_g_* = 8.0. Candidate feature pooling field centers were spaced 1.25 degrees apart (regardless of size) for a total of *G* = 2250 candidate feature pooling fields. The model for each voxel was run for 20 epochs of stochastic gradient descent with batch size of 200 and step size of 10^−4^ starting from an initial state of **w** = **0**.

#### 2.3.3. Layerwise Deepnet regression model (Deepnet-lReg)

In addition to the fwRF models described above we constructed a layerwise deepnet regression model for each voxel, as shown in Figure 2, following the work of Ref. [2]. For each voxel, 9 independent models (corresponding to the 5 convolutional layers, three fully-connected, and the final label probability layer of the DNN, respectively) were estimated and evaluated on a held-out subset (a randomly selected 10% of the samples) of the training data. Suppose that the model based upon layer *l* has the lowest loss on the held-out subset of the training data. If we denote pixel (*i, j*) of the *k^th^* feature map of layer *l* as 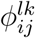, then the Deepnet-lReg model is specified as follows:

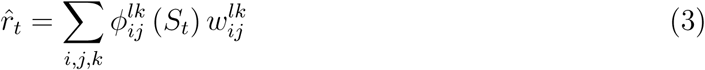

Note that this layerwise regression model assigns an independent weight to each pixel in each feature map, resulting in a total of K*_l_* × *N_l_* parameters, where *K_l_* and *N_l_* are the depth and number of units, respectively, of layer *l*. Weight optimization was performed via ridge-regression. The ridge hyperparameter was estimated using a brute force search over 14 log-spaced values between 10^−6^ and 10^8^. The layer/hyperparameter pair that minimized cost on the held-out subset was considered optimal and retained for further analysis (Fig.2).

### 2.4. Intepretation of the feature pooling field radius

The feature pooling field specifies the center and radius (i.e., standard deviation) of a 2D isotropic Gaussian function, *g*, that is applied to each feature map in the encoding model. It is important to emphasize that the pooling radius *σ_g_* models pooling over *feature map pixels*, not pixels in the stimulus. The pixels in any feature map will have their own intrinsic pooling radius, *σ*_f_, that specifies the region of the visual stimulus over which pixels are pooled to compute the feature transform (Fig 3). For some models the pooling radius of the feature map pixels is known explicitly. For example, in the population receptive field (pRF) model of Ref. [9], the pooling radius of each feature map pixel is determined solely by the downsampling applied (if any) to the visual stimulus as a pre-processing stage before model fitting. In the case of the Gabor-fwRF, the pooling radius of the feature map pixels is determined by the Gaussian envelope of the Gabor wavelet, which is known explicitly. However, note that the situation is slightly more complicated for the Gabor-fwRF model because the feature maps span multiple scales, so that pixels in feature maps associated with high-frequency wavelets have a smaller pooling radius than pixels in feature maps associated with low-frequency wavelets. The situation is even more complicated for the Deepnet-fwRF model. Not only do the feature maps of Deepnet-fwRF model scan multiple scales, but the exact pooling radius of the pixel in any feature map is only implicitly specified by the convolutional filters learned when the DNN was trained to label objects. Thus, although it is safe to assume that feature map pixels in the top layers of the DNN have a larger pooling radii than pixels in lower layers, the exact radii can be difficult to estimate. Given these considerations, the pooling radius in most fwRF models must be treated as a lower bound on the pooling radius that would be obtained if a pRF analysis were applied to data from a dedicated retinotopic mapping experiment. In what follows, we treat *σ*_f_ as the pooling radius of pixels in the feature map that makes the largest contribution to predicting voxel activity, and make use of the following approximation:

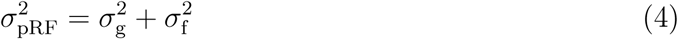
where *σ*_pRF_ is the standard deviation of the Gaussian pooling function used in a pRF analysis. For the Gabor-fwRF model (where *σ*_f_ is known) we apply this relation and report the pRF radius *σ*_pRF_. For the Deepnet-fwRF model, we will report only the feature pooling field radius *σ_g_*.

**Figure 3:**
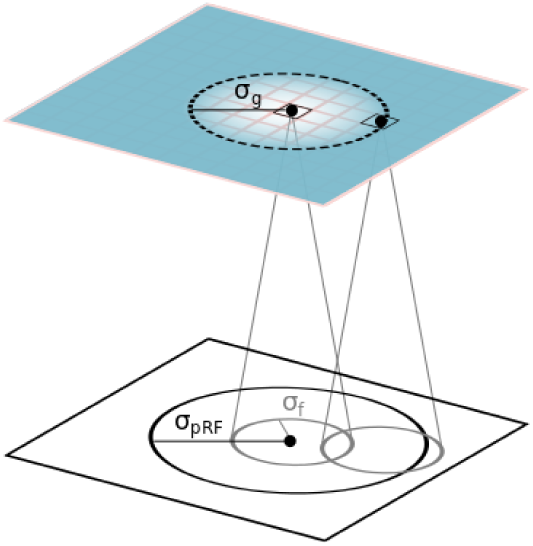
Interpretation of the feature pooling field radius. The feature pooling field in the fwRF model specifies the pooling radius (*σ*_g_) over feature map pixels (feature map illustrated as blue plane; pixels indicated by pink grid; feature pooling field illustrated as black dashed circle). The pixels in a feature map (two pixels are labeled by black dots) have their own intrinsic scale that is specified by a pooling radius (*σ*_f_, solid gray circles in white plane) over pixels in the stimulus (white plane). Assuming that both the feature pooling field and the pooling field of the feature map pixels are Gaussian, an estimate of the square of the population receptive field radius (σ_p_R_F_, large solid black circle in the white plane) can be obtained by summing the square of the feature pooling field radius with the square of the pooling radius of the feature map pixels.

### 2.5. Model evaluation comparison

For each voxel, the prediction accuracy of each of the three models was evaluated and compared. Model evaluation was performed by correlating a model’s predicted responses 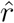 with measured responses *r* across all 120 samples in the validation set. This resulted in a measure of *prediction accuracy ρ* ∈ [−1, 1] for each voxel. To compare models, we constructed accuracy/advantage plots ( Fig. 7). To construct these plots, we first created a scatter plot in which each dot corresponds to a single voxel. The position of each dot along the vertical axis indicates the average prediction accuracy of the two models being compared, while the offset to the left or right of the vertical line at 0 indicates the difference in prediction accuracy between the two models. Thus, position along the horizontal axis indicates the advantage in prediction accuracy of one model relative to the other. Second, the individual voxels were binned to create an estimate of voxel density in the accuracy/advantage plane.

**Figure 4:**
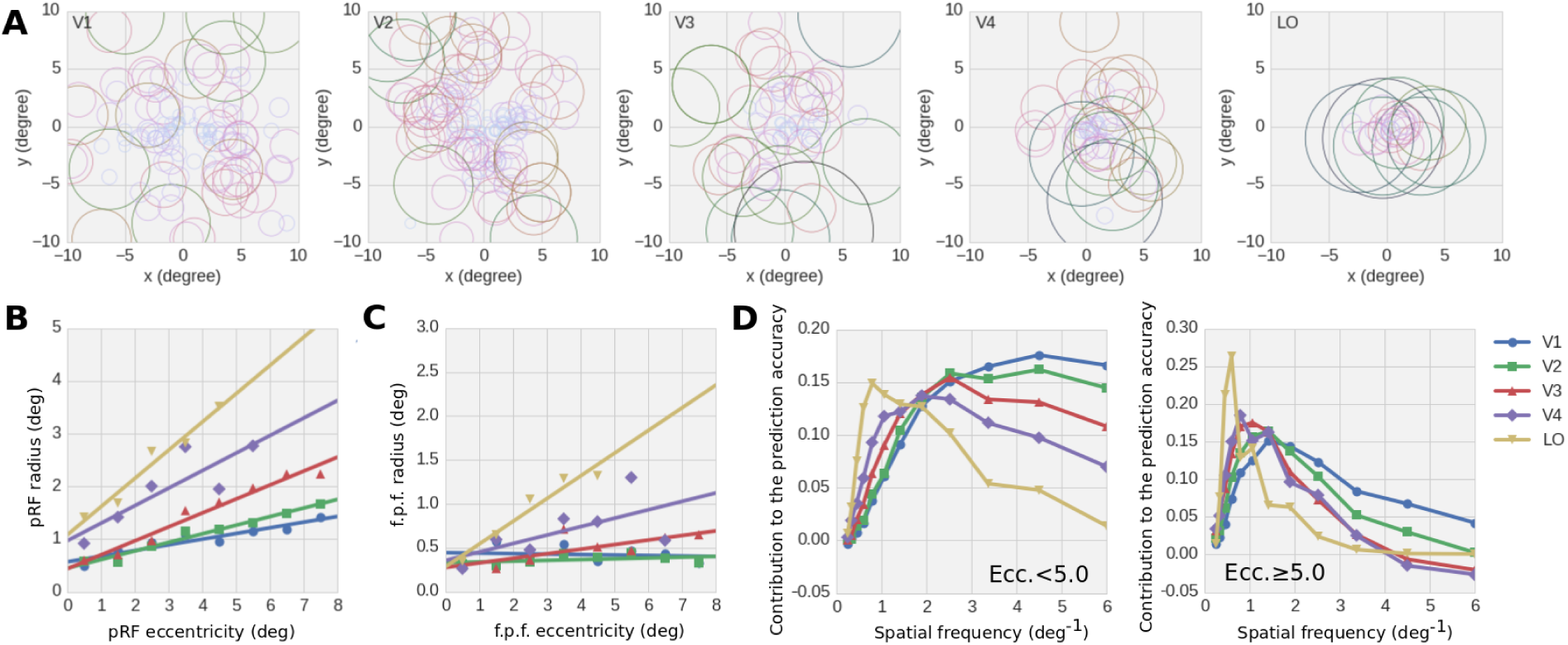
Feature pooling fields and spatial frequency tuning for the Gabor-fwRF model. (A) Panels show data from distinct ROIs (labeled in upper left corner). Each circle shows the feature pooling field of a single voxel (only voxels with *ρ* > 0.2 are included in these plots). The radius of each circle is the pRF radius (σ_p_R_F_) estimated from the feature pooling field radius (*σ*_f_) using the relation of Eq. 4. Circles are color-coded according to radius for ease of interpretation. In early visual areas, receptive fields estimated by the Gabor-fwRF tend to be relatively small and scattered across the visual field; in higher visual areas, they are relatively large and concentrated at the fovea. (B) The average estimated pRF (*σ*_p_R_F_) is plotted against feature pooling field eccentricity. Color indicates brain ROI. Lines show the best linear fits. Radius increases linearly with eccentricity in all ROIs. (C) The feature pooling field radii also exhibit linear scaling with eccentricity, although they underestimates the pRF radii in all ROIs. (D) Spatial frequency tuning. Curves show the average contribution to the total prediction accuracy of each spatial frequency. As expected, the average preferred spatial frequency shifts downward from perifoveal (left) to peripheral eccentricities (right), and from lower to higher visual brain areas.

**Figure 5:**
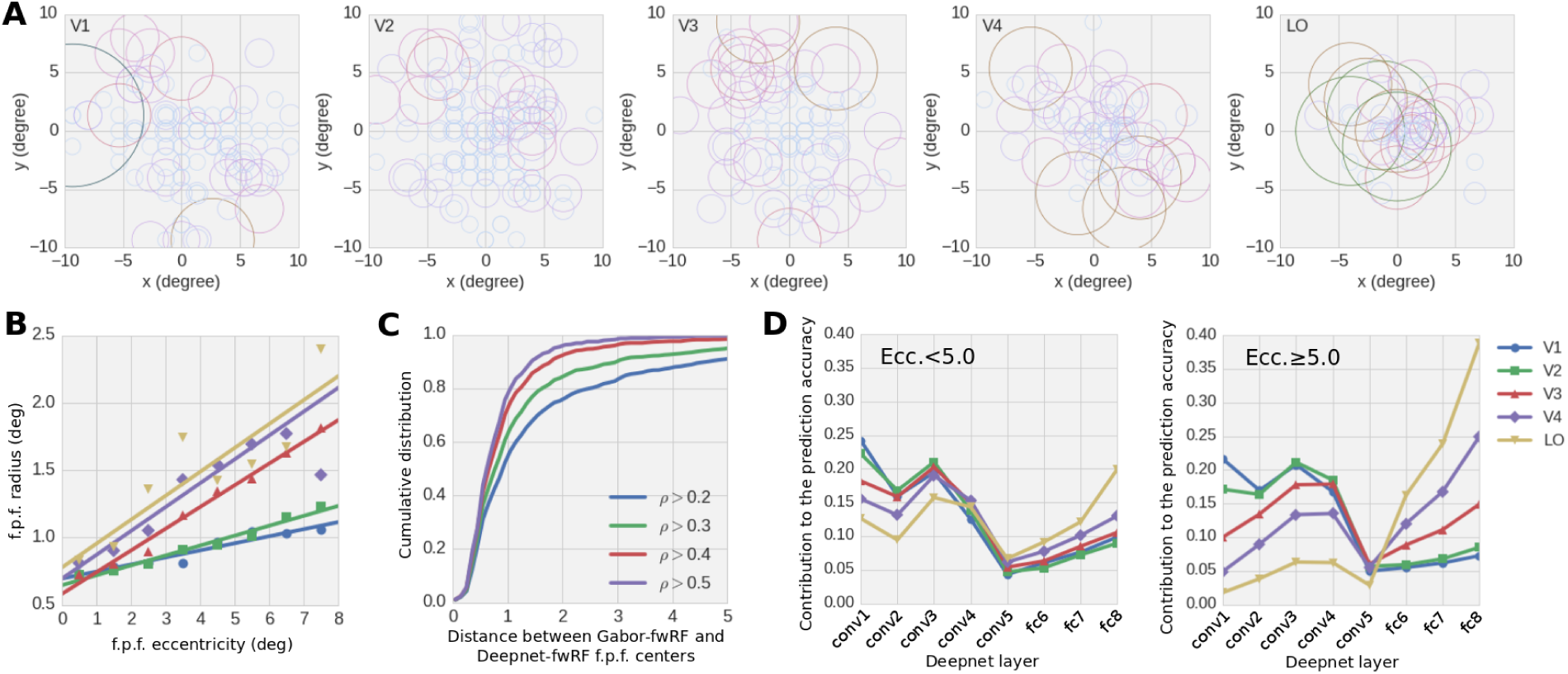
Feature receptive field and layer tuning for the Deepnet-fwRF. (A) Panels show data from distinct ROIs (labeled in upper left corner). Each circle shows the feature pooling field of a single voxel (only voxels with *ρ* > 0.2 are shown here). The radius of each circle is the feature pooling field radius (*σ_g_*). As in the Gabor-fwRF model, the feature pooling fields are relatively small and scattered for early visual areas, and relatively large and foveated for higher visual areas. (B) The Deepnet-fwRF uncovers a positive linear relationship between feature pooling field radius and center eccentricity. (C) Cumulative distribution of the distance between feature pooling field centers under the Deepnet-fwRF and Gabor-fwRF models. Each colored curve shows the cumulative distribution conditioned upon the prediction accuracy threshold given in the legend. Agreement between the models is generally high and increases with increasing predication accuracy threshold. (D) The average contribution to the total prediction accuracy of models in each brain area are plotted for each layer of the DNN for voxels with peri-foveal (left panel) and peripheral (right panel) feature pooling field centers. Models in all brain areas receive significant contributions from multiple DNN layers. The contribution of the early DNN layers is attenuated for higher visual cortical areas, while the reverse trend occurs for deep layers of the DNN hierarchy.

**Figure 6:**
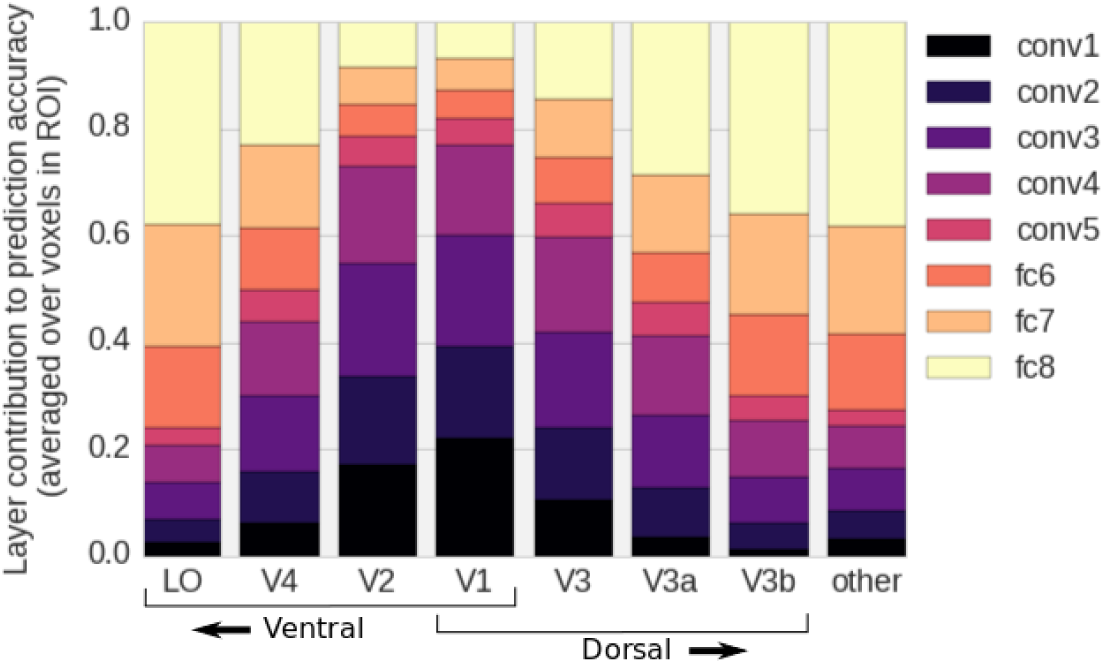
Contributions of the DNN layers to dorsal and ventral stream predictions. Each column shows the distribution of the DNN layer contributions to the prediction accuracy for a single ROI. Colored bars within each column indicate the contribution to the prediction accuracy averaged over all voxels in that ROI. The contribution of the lowest (conv1) and highest (fc8) layers exhibit a clear counter-gradient organization. Contributions of intermediate DNN layers are also graded, but are much more uniformly distributed across ROIs.

**Figure 7:**
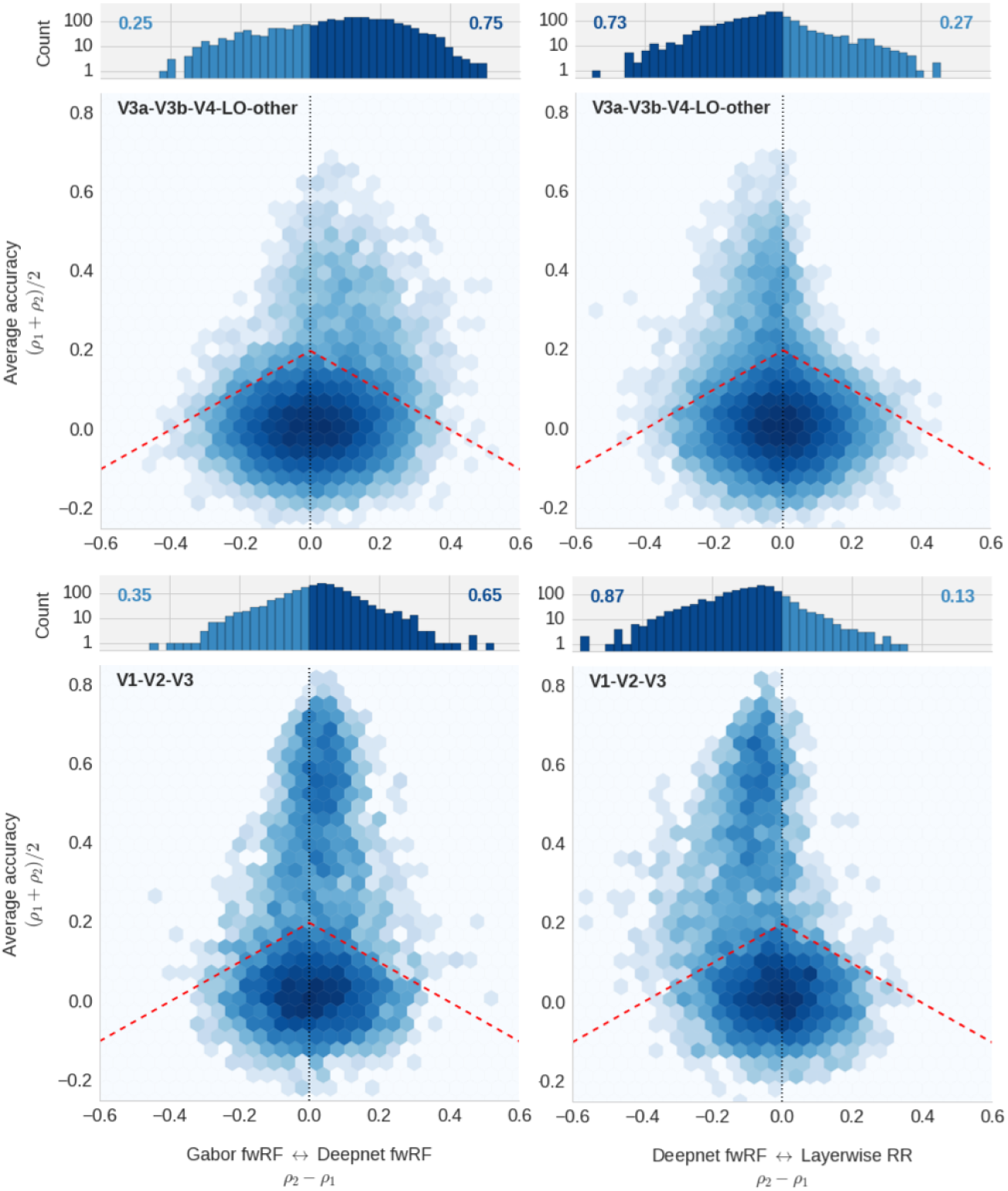
Comparison of the Gabor-fwRF, Deepnet-fwRF, and layerwise deepnet regression models. Each of the four accuracy/advantage plots displays a comparison of prediction accuracies for two models. The position along the vertical axis indicates the average prediction accuracy for the models under comparison; shifts to the right or left along the horizontal axis indicated a relative improvement in prediction accuracy for one of the models (model 1 is presented to the left of model 2). The color of each hexagonal bin indicates the number of voxels in a local region of the plot (log scaled). The histogram at the top of each plot represent the distribution of relative improvements for all voxels whose prediction accuracy is above 0.2 for at least one of the two models, which correspond graphically to all voxels above the red dashed line. The number on each side represents the fraction of voxels that are improved under that model. In the plots on the left, a shift in the data towards the left indicates an advantage for Gabor-fwRF model. In plots on the right, a shift of the data towards the right indicates an advantage for the Deepnet-lReg model. In all plots, a shift of the data toward the midline indicates an advantage for the Deepnet-fwRF. The upper plots display data for voxels in intermediate and higher visual areas (V4, V3A, V3B, LO, and “other”); the lower plots display data for voxels in the early visual cortex (V1, V2, V3). For intermediate brain areas, the Deepnet-fwRF outperforms both the layerwise regression and Gabor-fwRF models. For early visual areas, the Deepnet-fwRF strongly outperforms the layerwise regression model, but only weakly outperforms the Gabor-fwRF. The Deepnet-fwRF thus seems to have the strongest overall advantage for brain areas that require complex feature spaces. The “banana” shape of the distribution in the lower right suggests that the fwRF model provides strong and appropriate regularization, since voxels with low prediction accuracy under the more complex layerwise regression model are effectively “rescued” by the Deepnet-fwRF.

### 2.6. Feature map contribution to the prediction accuracy

For both the Gabor-fwRF and the Deepnet-fwRF, the feature maps all contribute linearly to the final prediction according to Eq. 1. While it may be common to look at the weights of this linear combination to determine the relative importance of each contribution, the value of the weights themselves are dependent upon the typical values of each feature map, such that it makes any comparison across feature maps difficult. For this reason, and due to the linearity of the model, we chose to look instead at 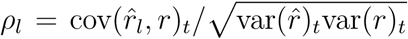 where 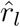 is the same as Eq. 1 but where the index *k* over feature maps only runs over *k* ∈ *K_l_*, a subset of feature maps sharing certain properties. For a list of disjoint subset K*l* that cover all *K* feature maps, it follows that *∑_l_ρ_l_* = *ρ* where *ρ* is the Pearson correlation coefficient between the actual and predicted activity for that voxel. We therefore call these *ρ_l_*’sthe contributions to the total prediction accuracy.

## 3. Results

We fit and then evaluated fwRF models using a previously published and publicly available dataset [6, 7]. The dataset contains estimates of functional BOLD activity in response to greyscale natural photographs from voxels in visual brain areas V1, V2, V3, V4, V3A, V3B, and LO. Voxels in visually responsive cortex anterior to LO, labeled “anterior occipital cortex” in a previous publication [7] are also included.

### 3.1. fwRF models recover feature pooling fields and tuning functions for both simple and complex features

An established encoding model for early visual areas is the Gabor Wavelet Pyramid (GWP) model [6, 7]. To ensure that the fwRF modeling approach is compatible with this established model we first designed a fwRF version of the GWP, referred to here as the Gabor-fwRF model. To construct feature maps for the Gabor-fwRF model each photograph in the experimental stimulus set was convolved with each of 96 complex Gabor wavelets (12 spatial frequencies, 8 orientations) and then passed through an elementwise nonlinearity (see Methods for complete details). This procedure produced 96 distinct feature maps that were used for the Gabor-fwRF model. For each voxel in the dataset we estimated the optimal feature weights and feature pooling field by performing gradient descent on the feature weights and brute-force grid search on the feature pooling field radii and centers. The Gabor-fwRF feature pooling fields conformed to well-known patterns of visual receptive field organization (Fig. 4A-C). In lower visual areas, feature pooling fields are relatively small and uniformly tile the visual field (e.g., Fig. 4A, V1 panel) while in higher visual areas, receptive fields are relatively large and concentrated at the fovea (e.g., Fig. 4A, LO panel). In all areas, the Gabor-fwRF uncovers a positive linear relationship between pooling size and eccentricity (Fig. 4B). The slope of the lines relating the feature pooling field eccentricity to the estimated pRF radii *σ*_pRF_ in each ROI are comparable to those presented in previous studies [9]. As expected, feature pooling field radii (*σ_g_*) bound the pRF radii from below (Fig. 4C), and, like the pRF, exhibit an attenuated size/eccentricity relationship in early visual areas.

The spatial frequency tuning functions of the Gabor-fwRF model (Fig. 4D) also conform to well-known properties of visual cortical organization. Voxels with feature pooling field centers that are relatively close to the fovea prefer (on average) relatively high spatial frequencies. Lower visual areas prefer larger spatial frequencies more than higher visual areas. Voxels with receptive fields relatively far from the fovea prefer relatively small spatial frequencies, with lower visual areas again having a higher spatial frequency preference than higher visual areas.

A known failing of encoding models that rely on Gabor-like visual features is that their prediction accuracy becomes increasingly poor when applied to intermediate and higher cortical visual areas. However, recent work [2, 4] has shown that feature maps sourced by DNNs that have been trained to classify objects can be used to significantly improve the prediction of encoding models for intermediate and high-level visual areas. We therefore used the fwRF modeling approach to design a Deepnet-fwRF in which the feature maps are taken from the internal representations of a deep convolutional neural network. The Deepnet-fwRF model included all feature maps from 5 distinct convolutional layers, as well as up to 1, 024 feature maps per layer from 3 fully-connected layers. This resulted in a fwRF model with 4, 424 feature weights.

The Deepnet-fwRF model reveals retinotopic organization consistent with that revealed by the Gabor-fwRF model. The feature pooling field radius recovered from the models exhibit a positive linear dependence upon eccentricity, and increase monotonically across the hierarchy of visual ROIs. Voxelwise estimates of feature pooling field center were consistent with estimates obtained from the Gabor-fwRF model Figure 5C. The distance between centers provided by the two models decreased with the quality of their predictions (up to a limit due to the finite grid of feature pooling field centers used by the optimization algorithm). These results suggests that the ability of the fwRF modeling approach to uncover the retinotopic organization of visual areas is relatively insensitive to the feature maps used in the encoding model, so long as those feature maps confer accurate model prediction accuracy.

As expected, the Deepnet-fwRF also uncovered feature tuning functions that align the increasing complexity of the network’s feature maps with the increasing complexity of visual representations in the brain (Fig. 5D). The average contribution of feature maps in each layer of the network to each ROI depended on the position of the ROI within the visual hierarchy. Deepnet-fwRF models for V1 and V2 assigned little contribution to network layers 5 and higher while the contributions assigned to layers 5 and higher increased dramatically for later visual areas V3, V4, and LO. However, note that even ROIs that received a relatively strong contribution from a single layer (e.g., V1 and DNN layer “conv1”) nevertheless received a non-negligible contribution from other DNN layers as well (Fig. 6). This is especially true for voxels with peripheral feature pooling field centers Fig. 5D, right panel). For these voxels the Deepnet-fwRF model distributes contributions across DNN layers much more uniformly. This suggests that the Deepnet-fwRF’s ability to linearly combine feature maps across layers is a potentially important attribute of the model.

### 3.2. The Deepnet-fwRF model predicts activity more accurately than less expressive models

Our analysis of the Deepnet-fwRF feature weights demonstrate that feature maps from across the DNN hierarchy contribute to prediction accuracy in all brain ROIs (Figure 6). The capacity for cross-layer blending of feature maps makes the Deepnet-fwRF model highlyexpressive, which may in turn allow it to make more accurate predictions than models that are less expressive. To test this hypothesis, we compared the prediction accuracy of the Deepnet-fwRF to the Gabor-fwRF model and to an encoding model that runs a layerwise regression of DNN feature maps (Deepnet-lReg). Under the Deepnet-lReg model, weights are assigned to every pixel in every feature map of a single DNN layer that is selected via a brute-force optimization procedure. Thus the Deepnet-lReg model effectively imposes a hard layerwise sparseness constraint. This can be contrasted to fwRF models, which impose a hard spatial constraint in the form of space-feature separability, but do not impose any constraint on the combination of DNN layers that contribute to each model.

We found that, for the current dataset, the Deepnet-fwRF model has a significant advantage in prediction accuracy over both the Gabor-fwRF and Deepnet-lReg models (Fig. 7). For visual areas V1, V2, and V3 the advantage over the Gabor-fwRF model is subtle in general and is in fact non-existent for voxels with feature pooling field centers near the fovea (Fig. 8). The advantage of the Deepnet-fwRF model over the Deepnet-lReg model in these areas is more pronounced. The Deepnet-fwRF model more accurately predicts activity for 87% of these voxels, including voxels for which the Deepnet-lReg model fails completely. For visual areas V4, V3A/B, and AOC the Deepnet-fwRF model also enjoys an advantage in prediction accuracy, out-predicting the Gabor-fwRF and Deepnet-lReg models for an average of 74% of the voxels.

**Figure 8:**
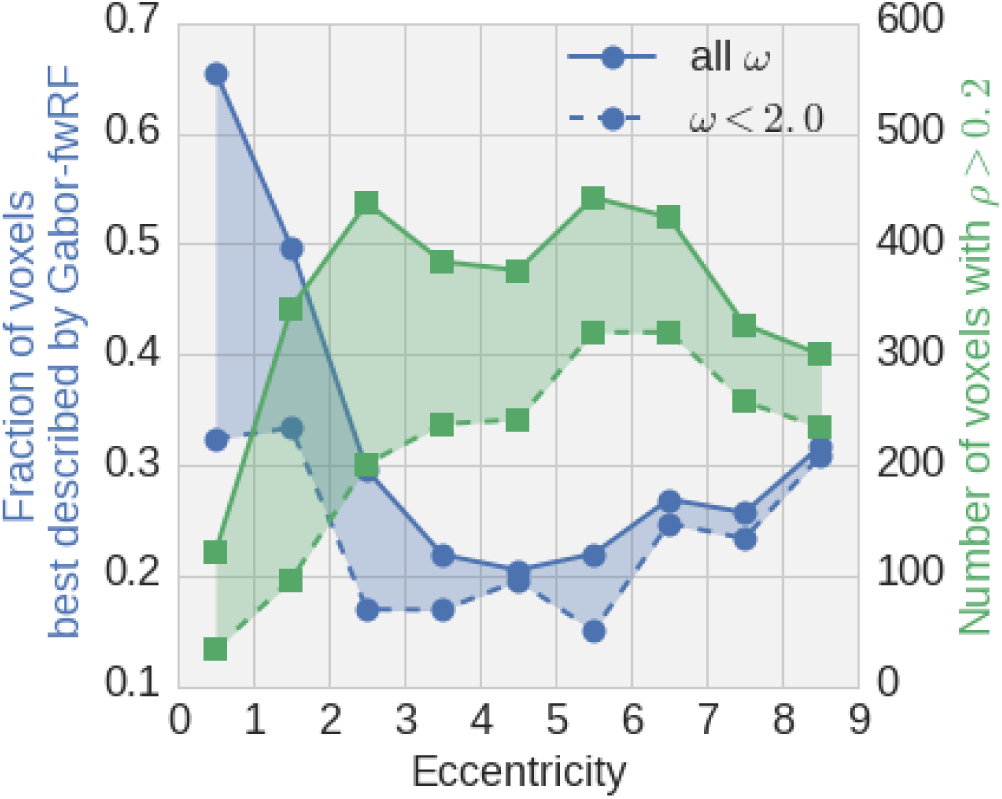
Model advantage as a function of eccentricity. The blue curves indicate the fraction of voxels (left vertical axis) for which the Gabor-fwRF model has higher prediction accuracy than the Deepnet-fwRF model as a function of eccentricity (average feature pooling field center eccentricity from Gabor-fwRF and Deepnet-fwRF models; horizontal axis). The green curves indicate the number of voxels (right vertical axis) available for analysis at each eccentricity, with a bin width of 1 degree. The Gabor-fwRF model performs better than the Deepnet-fwRF model for foveal voxels that prefer high spatial frequency. The advantage of the Gabor-fwRF for very foveal voxels disappears when the analysis is restricted to voxels with low spatial frequency preference (*ω* < 2 cycles/deg.; dashed curves).

## 4. Discussion

We have introduced the feature-weighted receptive field (fwRF), a new approach to building voxelwise encoding models for visual brain areas. The results of this study suggest that the fwRF modeling approach can be used to achieve the performance goals of expressiveness, scalability, interpretability and compatibility laid out above. The key design principle of the fwRF modeling approach is space-feature separability, which endows the model with an explicit receptive field-like component that facilitates interpretation, and makes it possible to scale the number of feature maps in the model without incurring a per-pixel increase in model parameters. We find that when this approach is applied to a deep neural network with thousands of feature maps, the resulting encoding model achieves better prediction accuracy than comparable encoding models for most voxels in the visual system.

### 4.1. Relationship to previous work

Several important voxelwise modeling approaches have preceded the fwRF modeling approach. One very popular and important class of models are those designed specifically for retinotopic modeling. This includes the inverse retinotopy method [15] and the population receptive field method [9]. These methods permit estimation and detailed analysis of the locations and sizes of voxelwise receptive fields. However, they do not include an explicit feature map. As a consequence, they do not reveal any information about tuning to visual features (e.g., spatial frequency, orientation) and must be estimated with dedicated retinotopic mapping experiments that utilize stimuli optimized for the purpose.

The fwRF model is a special case of the linearized, regularized regression approach described in [16]. This more general approach also depends upon the construction of a set of nonlinear features. Unlike the fwRF, these nonlinear features are not required to be arranged in a feature map. Like the fwRF, they represent hypotheses about the visual features encoded in brain activity. The fwRF model overcomes two limitations of this more general approach. First, in the more general approach the number of model parameters scales with the number of feature map pixels. This pixelwise scaling necessarily imposes a trade-off between the resolution and the number of feature maps used in the model. Second, under the general regression approach deriving an explicit receptive field and feature tuning function from models weights often requires in-silico experiments on the models. This can in fact be a very powerful tool for model interpretation, but ideally it should not be necessary for recovering basic receptive fields and feature tuning properties.

### 4.2. Using the fwRF to measure receptive field size

The fwRF model yields a receptive field-like size measure that we have referred to as the “feature pooling field radius”. We took pains to emphasize that this measure, *σ*_g_, is importantly different from the more familiar population receptive field (pRF) size, *σ*_pRF_, which is in turn importantly different from the classical receptive field sizes of single neurons [9]. The differences between these measures underscore the fact that receptive field size depends entirely upon the features being pooled over. Thus whenever possible pooling sizes should be interpreted in light of the pooling required to compute the features being represented.

This “feature pooling”, as we have called it, is not always easy to estimate. However, we have shown that even when these estimates are not available, the feature pooling field radius *σ*_g_ can provide a lower bound on pRF size that exhibits the positive relationship between size and eccentricity that is a hallmark of visual cortical organization.

### 4.3. Using the Deepnet-fwRF to map changes in representational complexity across visual areas

Our Deepnet-fwRF model results are consistent with those reported in Ref. [2], which show that changes in representational complexity across visual brain areas and layers of the DNN are well-aligned. Nonetheless, analysis of the Deepnet-fwRF model weights (Figures 5D and 6) shows that the contributions to the predictions for any ROI are widely distributed across DNN layers, particularly for voxels with peripheral feature pooling field centers. This may reflect the fact that activation in different DNN layers can make redundant contributions to the Deepnet-fwRF model predictions. For example, “conv1” alone may be able to explain 90% of the explainable variance of a voxel in V1 while, at the same time, “conv2” alone would be able to explain 85% of it, with a large overlap between the two predictions. As a result, the variance explained by the Deepnet-fwRF model is distributed across layers when several layers are used in conjunction. However, some explained variance remains unique to specific layers which account for the clear gradient of complexity observed.

Our analysis also shows that idiosyncrasies of the underlying DNN can violate the regularity of the alignment of ROIs and DNN layers, with some layers contributing only very little in all ROIs. For example, the highest convolutional layer “conv5” made the smallest contribution to all ROIs, which occurs near the inflection point in the layer tuning functions for all brain areas (Fig. 5D). This could be related to the fact that “conv5” is itself a special point in the network architecture where the network switches from a convolutional to a fully-connected architecture after “conv5”.

### 4.4. Prediction accuracy advantage of the Deepnet-fwRF

The general advantage in prediction accuracy of the Deepnet-fwRF model over the Gabor-fwRF and Deepnet-lReg models is a strong endorsement for the fwRF modeling approach. Our results suggest that the Deepnet-fwRF out-predicts the Gabor-fwRF model because the Deepnet-fwRF contains feature maps that are more appropriate for explaining intermediate visual areas. While effective models for early visual areas based on Gabor-like features, and higher visual areas based on semantic features [7, 10, 17], have been available for some time, intermediate visual areas have been most resistant to modeling. The delivery of an encoding model that makes predictions for intermediate areas that are as accurate as those of the aforementioned models for early and object-specific visual cortex would seem to be one of the most salient findings. Exactly what contribution the intermediate visual areas make to visual processing is still unknown, since the function of the DNN layers that most strongly contribute to these areas in the Deepnet-fwRF model is unknown. However, these prediction results effectively quarantine the problem, replacing the challenge of interrogating the brain *in vivo* with the challenge of interrogating a DNN network *in silico*.

There are three possible reasons for why the Deepnet-fwRF model out-predicts the Deepnet-lReg model, none of which are exclusive. One possible reason is the increased expressiveness of Deepnet-fwRF model. Analysis of Deepnet-fwRF model weights suggest that the model takes advantage of this expressiveness by distributing weights across layers. A second, related possibility is that the space-feature separability imposed by the Deepnet-fwRF model is a more appropriate form of regularization than the layerwise sparseness (and within-layer smoothness) imposed by the Deepnet-lReg model. Supporting this possibility is the fact that the Deepnet-fwRF model seems to be able to accurately predict activity in voxels that are predicted very poorly by the Deepnet-lReg model, suggesting that the Deepnet-fwRF model more effectively rescues voxels with poor signal-to-noise characteristics. Finally, it is well-known that a model’s prediction accuracy depends heavily upon an accumulation of many idiosyncratic choices. An alternate set of equally reasonable choices might have produced a less stark contrast between the prediction accuracy of the Deepnet-fwRF and Deepnet-lReg models. We therefore interpret these results as a suggestion that the increased expressiveness made possible by the Deepnet-fwRF model is a worthwhile attribute that merits future application and further experimentation.

### 4.5. Dynamic fwRF models

The fwRF models in this study are static in the sense that predictions of activity at time *t* depend only on concurrently presented stimuli, and not on the past history of stimulus presentation. This choice was appropriate because the temporal dynamics of the voxel activities had already been modeled out of the data (see Ref. [6]). However, the fwRF approach can easily accommodate dynamic datasets by including time-shifted copies of each feature map, such that

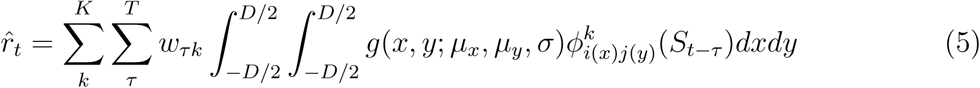
where τ ∈ [0, *T*] indexes time shifts. Under this approach, each feature map would have an associated temporal kernel ***w**_k_* = (*w*_0*k*_, …, *w_Tk_*) instead of a single static weight *w_k_*. These temporal kernels would be estimated via gradient descent.

Alternatively, we might enforce space-*time*-feature separability by including an explicit temporal kernel function

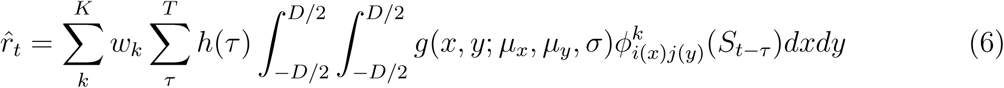
where *h*(·) is the explicit temporal kernel function. Ideally, this function would have a small number of shape parameters that would be estimated–like the receptive field parameters–via brute-force search.

### 4.6. The Gaussian pooling field

The fwRF models presented here used a 2D Gaussian feature pooling field whose radius was fixed for all feature maps. While keeping the radius constant has the advantage of reducing the number of parameters of the model, it also reduces its expressiveness. Allowing the feature pooling field radii to vary across the feature map would allow the model to capture, among other things, receptive fields with a “Mexican hat” profile that enforce a suppressive band around an excitatory center. Furthermore, any receptive field function with a small number of shape parameters could be trained using the gradient-descent with grid-search optimization algorithm presented here. There may very well be a more optimal choice of feature pooling field structure, but we will need future studies to confirm this.

## 5. Conclusions

We have introduced the feature-weighted receptive field (fwRF), a new approach to building voxelwise encoding models for visual brain areas. The results of this study suggest that the fwRF modeling approach has satisfied its four stated performance goals of expressiveness, scalability, interpretability and compatibility. The key design principle of the fwRF modeling approach is space-feature separability which makes it possible to consider large number of feature maps without incurring a per-pixel increase in model parameters. Finally, when applied to a deep neural network with thousands of feature maps, the resulting encoding model achieved state-of-the-art prediction accuracy for voxels in the visual system.

## Acknowledgements

This work was supported by grant NIH R01 EY023384 to TN.

## A Appendix

### Hardware

All instances of the fwRF model presented in this paper and the model throughput benchmark have been run on a system equipped with a Intel 6 cores i7-5930K processor, 128 Gb of RAM, and a dedicated NVIDIA Titan X (Maxwell) video card (12 Gb of VRAM).

### Description of the fwRF algorithm

The fwRF algorithm is divided into two main procedures: First, we calculate a *model-space* tensor, which is the most memory intensive part of the algorithm. Second, we learn the model weights through (stochastic) gradient descent and infer the best parameters for the pooling function through brute force search. The main stokes of the algorithm are described in Algorithm A.1.

### Scaling of the current fwRF implementation

A short inspection of Algorithm A.1 would convince someone that the current fwRF implementation scales linearly in the number of voxels, number of candidate pooling function model and number of samples in the time series. The scaling with respect to the feature map size and the dependence on the number of feature maps is more difficult to assert. Since the feature map size only appears within the model space tensor evaluation due to our assumption of space-feature separability, and that this operation was usually much faster than the model weight estimation (it remained on the order of a few minutes even with feature maps sizes of a few hundred pixels), the only factor relevant to the model performance is the number of feature maps. Figure A.1 shows the typical model throughput as a function of the number of feature maps (assuming adequate choices of batch size across voxel, candidate models and samples). The batch sizes were selected to reach optimal utilization of the GPU resources.

#### Algorithm A.1

Feature weighted receptive field method. In the following, tensors are boldfaced and the dimensions of a tensor are always displayed in square bracket after its name. A change in the order of the dimension implies a transpose of the tensor and an arrow in the dimensions indicates broadcasting to a certain size.

~~~
1: **procedure** Generate model space tensor(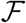, 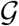):
2: **for** (**F***_l_*, g*_l_*) ∈ (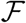, 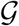) **do**
3: **M***_l_* ← TensorDot(**F***_l_*[*n, K_l_, x_l_, y_l_*], *g_l_*[*G, x_l_, y_l_*], axis = [[2, 3], [1, 2]])
4: **M** ← Concatenate((Ml[*n, K_l_, G*], ∀*l* ∈ 2 L), axis = 1)
5: **M** ← Z-Score(**M**, axis = 0)
6: **return M**[*n, K, G*]
7:
8: **procedure** Generate voxel prediction(**M**,**w**):
9: 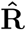 ← Batched-TensorDot(**M**[*G, n, K*], **w**[*G, V, K*], axis = [[2], [2]])
10: return 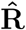[*n, V, G*]
11:
12: **procedure** LOSS(**M**, **R**, **w**):
13: 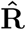 ← Generate voxel prediction (**M**, **w**)
14: **return** L2-norm(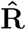[*n, V, G*] − **R**[*n, V*, 1 → *G*])[*V, G*]
15:
16: **procedure** Optimize FWRF model parameters(**M**,**R**,**w**_init_):
17: **m**_best_ ← **zeros**[*V*] # best models
18: **w**_best_ ← **zeros**[*V,K*] # best weights
19: **s**_best_ ← **inf**[*V*] # best scores
20: **for** *V_b_* ∈ Batch(*V*) **do**
21: **for** *G_b_* ∈ Batch(*G*) **do**
22: **w***_b_* ← **w**_init_[1 → *G_b_*,*V_b_, K*]
23: **s***_b_* ← **zeros**[*V_b_, G_b_*]
24: **for** *e* ∈ 1˙.epochs **do**
25: **for** *n_b_* ∈ Batch(*n*_train_) do
26: **w***_b_* ← **w***_b_* − λ 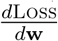 (**M**[*n_b_, K, G_b_*], **R**[*n_b_, V_b_*],***w**_b_*[*G_b_, V_b_, K*])
27: **for** *n_b_* ∈ Batch(*n*_holdout_) **do**
28: ***s**_b_* ← ***s**_b_* + Loss(**M**[*n_b_, K, G_b_*], **R**[*n_b_, V_b_*], *w_b_*[*G_b_, V_b_, K*])
29:
30: **for***υ* ∈ Element(*V_b_*_)_**do**
31: *g* ← argmin(*s_b_*[*υ, G_b_*])
32: if *s_b_*[*υ, g*] < **s**_best_[*υ*] **then**
33: **m**_best_[*υ*] ← *g*
34: **w**_best_[*υ, K*] ← **w**_*b*_[*g, υ, K*]
35: **s**_best_[*υ*] ← **s**_*b*_[*υ, g]*
36: **return m**_best_, **w**_best_, **s**_best_
~~~

**Figure A.1:**
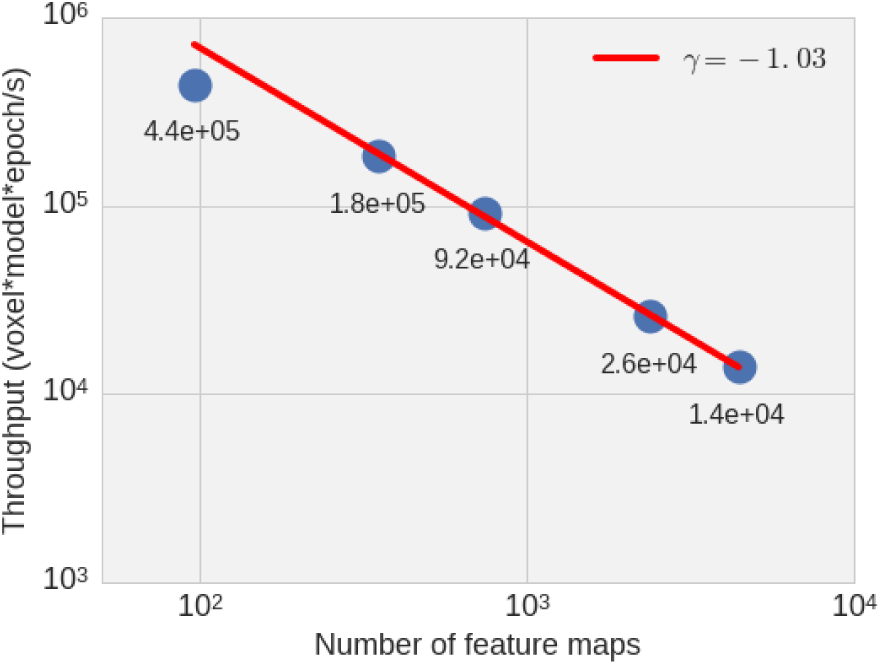
Scaling of the fwRF throughput. The throughput is expressed in voxel-model-epoch (vme) per seconds. The main factor determining the throughput is the number of feature maps, as shown here, which has almost optimal inverse scaling in a regime of full utilization. The batch size across voxels, model candidates and samples also affect the throughput, and the values displayed are for a fixed voxel batch size of 300 and a candidate batch size of 225. Overall computation time scales linearly with the sample size, number of voxels and number of candidates. For example, a case with 22K voxels and 20K candidate models optimized over 20 epochs results in 8.8 × 10^9^ vme. For 96 feature maps at throughput of roughly 5 × 10^5^ vme/s results in an estimated computation time of 4.9 hours. This does not account for the time required to prepare the model-space tensor, which is however usually much shorter than that.

